# The neurofunctional basis of affective startle modulation in humans – evidence from combined facial EMG-fMRI

**DOI:** 10.1101/567032

**Authors:** Manuel Kuhn, Julia Wendt, Rachel Sjouwerman, Christian Büchel, Alfons Hamm, Tina B. Lonsdorf

## Abstract

The startle reflex, a protective response elicited by an immediate, unexpected sensory event, is *potentiated* when evoked during threat and *inhibited* during safety. In contrast to skin conductance responses or pupil dilation, modulation of the startle reflex is valence-specific and considered *the* cross-species translational tool for defensive responding.

Rodent models implicate a modulatory pathway centering on the brainstem (i.e., nucleus reticularis pontis caudalis, PnC) and the centromedial amygdala (CeM) as key hubs for flexibly integrating valence information into differential startle magnitude.

We employed innovative combined EMG-fMRI measurements in two independent experiments and samples and provide converging evidence for the involvement of these key regions in the modulatory acoustic startle reflex pathway in humans. Furthermore, we provide the crucial direct link between EMG startle eye-blink magnitude and neural response strength.

We argue that startle-evoked amygdala responding and its affective modulation may hold promise as an important novel tool for affective neuroscience.

## Introduction

Defensive responding is innate and conserved across species with rapid protective reflexes promoting survival. However, ever-changing environments require flexible adaption ^1^. The mammalian startle reflex is elicited by an unexpected and abruptly occurring sensory stimulus (e.g. acoustic, tactile or visual) and is a prime example for the integration of short-latency responding and flexible modulation ^2,3^. It is conveyed through a sparse amount of synapses ^4–6^leading to a fast adaptation of eyelid-closure and body posture to prevent major injuries.

In humans, the startle eye-blink reflex represents the first and most reliable component of defensive responding ^7,8^. Importantly, this responding is modulated in a valence-specific manner (‘*affective startle modulation*, **ASM** ^9^): decreased (*inhibited*) during positive emotional states and increased (*potentiated*) during negative emotional states ^10^, such as when anticipating a potential threat (‘*fear potentiated startle*’, **FPS** ^3,11,12^). Due to this valence-specificity, startle responding represents an ideal tool for affective neuroscience - in particular as compared to other non-valence specific but commonly employed measures such as skin conductance responding (SCR) or pupil dilation ^13^.

To date, neurobiological models underlying this valence-dependent startle modulation are primarily derived from FPS studies in rodents ^5,6^and converge in implicating two distinct neural pathways: First, the *primary* acoustic startle pathway, conveying the startle response itself. Second, the *modulatory* pathway, adjusting response strength of the primary pathway depending on the current affective state - despite physically identical sensory input eliciting the startle response.

In rodents, the rapid *primary*acoustic startle reflex pathway involves three major hubs transferring the acoustic sensory input from the cochlear root neurons (CRNs) via the brainstem (i.e., nucleus reticularis pontis caudalis, PnC) to the motor-effectors that initiate the startle response ^4,14^.

The *modulatory*pathway, which is the focus of this work, centers on the pivotal role of the PnC as the key input hub for the integration of affective modulatory information. This modulatory input to the PnC is primarily conveyed through the centromedial nucleus of the amygdala (CeM) ^5,15–17^which is the core output region initiating defensive responding ^18,19^. Fine-tuning of this modulatory input is conveyed by regions exerting their influence either by modulating central amygdala activation or by direct input to the PnC (most prominently basolateral nucleus of the amygdala, BLA; bed nucleus of the stria terminalis, BNST; periaqueductal grey, PAG) ^5,20,21^.

The delineation of these neurobiological pathways has been exclusively derived from animal research but yet presumed to be universal across species. In humans, however, startle responding, in the fMRI environment, has been employed as an additional outcome measure of *emotional processing* ^22–26^while the neurobiological pathway underlying affective *startle responding* itself has not been investigated. Yet, startle responding is promoted as the ideal cross-species readout for fear and anxiety-related disorders ^27^, although evidence in humans is restricted to lesion ^28–30^and early PET imaging studies ^31,32^.

Based on the work in rodents, we here aim to delineate the neuro-functional basis of modulatory startle responding in humans, focusing on both the PnC and the CeM as key structures. More precisely, while rodent work is primarily based on fear-potentiated startle, we aim to demonstrate convergence and generality of this pathway across two well-established experimental approaches in humans: affective startle modulation (ASM) and fear-potentiated startle (FPS). Furthermore and importantly, we aim to provide a yet unexplored direct link between this defensive motor behavior (i.e., startle eye-blink magnitude) and neural activation to physically identical acoustic startle probes across emotional conditions in humans. This would further underscore the utility of startle responding as a unique tool for affective neuroscience.

To achieve these aims, we conducted two independent studies (ASMN=43, FPSN=55) and utilized recent methodological and technical advances: First we combined the acquisition of eye-blink startle and BOLD responding as assessed via facial electromyography (EMG) and functional magnetic resonance imaging (fMRI) respectively ^24–26^. Second, we utilized high resolution amygdala imaging as well as recent advances in human brainstem fMRI acquisition and data analysis ^33,34^.

## Results

### Identification of brainstem nuclei involvement in the primary acoustic startle reflex

To investigate the neural basis of affective modulation of the acoustic startle reflex it is essential to first delineate the neural basis of the *primary*acoustic startle pathway – investigated here by utilizing the startle habituation phase which involves repetitive startle probe presentations *without* emotional forground information in the ASM study (**Figure 1B**).

**Figure 1.**
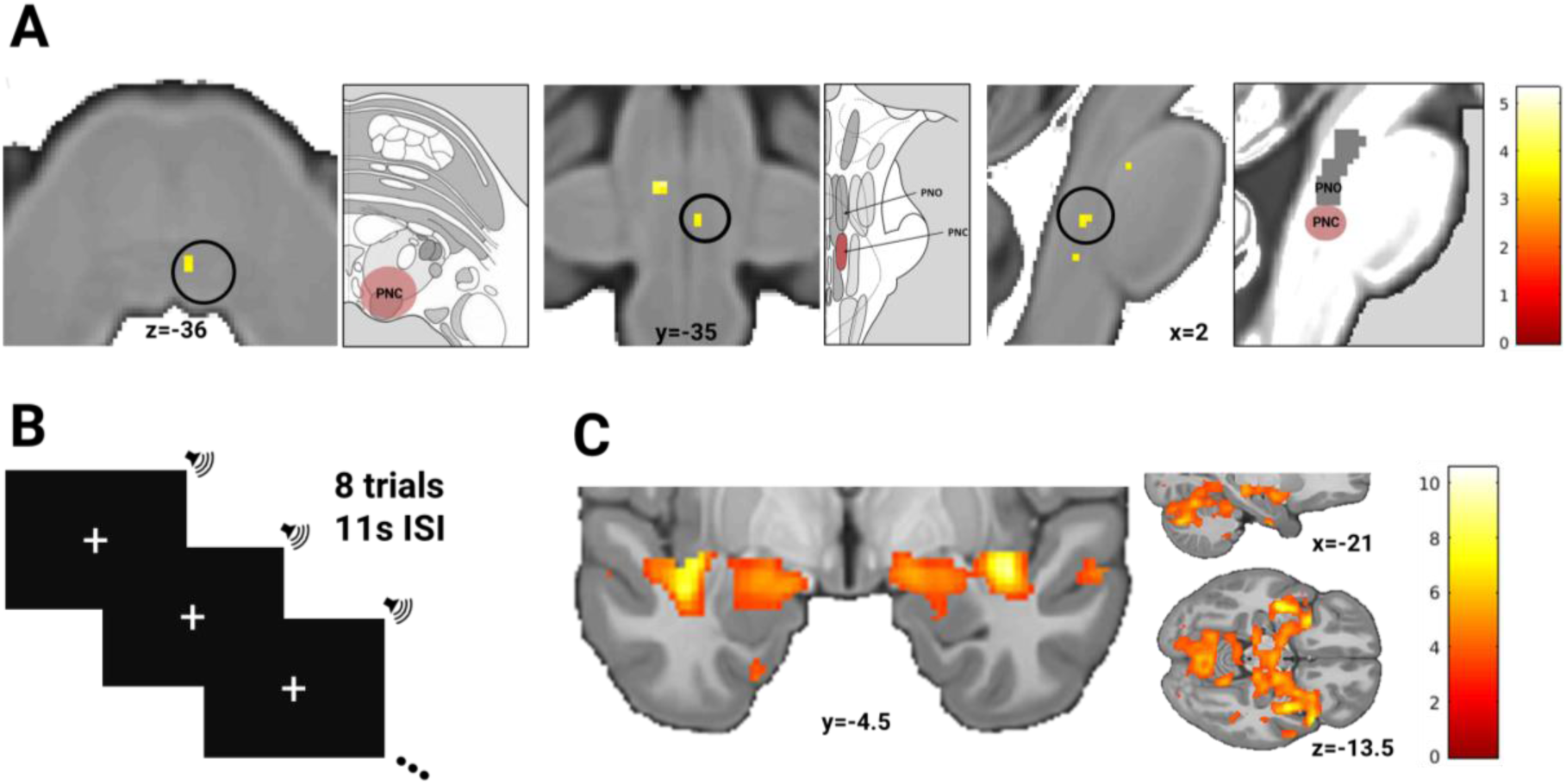
(A) Nucleus reticularis pontis caudalis (PnC) responding evoked by (B) eight acoustic startle probe presentations, suggesting PnC involvment in the primary acoustic startle pathway. (C) Concomittant responses towards the startle probe in the centro-medial amaygdala. Schematic illustrations in grey Boxes in A: Location of the PnC (highlighted in red) and the nucleus pontis caudalis oralis (PnO) as defined by Duvernoy’s Atlas of the Human Brain Stem and Cerebellum (left and middle, Naidich et al., 2009, adapted by permission from Springer Nature) as well as in reference to an available anatomically defined MRI ROI of the PnO (right, Edlow et al., 2012). Note that black circles highlight the activation within the PnC region and do not illustratethe specific size of search volume. Display threshold at p_uc_<0.001.

As expected from rodent work, we indeed observed PnC activation (p_uc_ < 0.001, T = 3.47, k = 3, [x,y,z] = [2,-35,-36], **Figure 1A, Table S1**) as well as concomittant activation in secondary ROIs (i.e., CeM, PAG, both pFWESVC<0.004, **Figure 1C; Table S1)** in response to repetitive startle probe presentation. This supports the proposed role of the PnC as key hub in the human *primary acoustic startle reflex* pathway (see SI for additional brain-behavior correlation), which sets the stage for investigating the involvement of the proposed core regions (i.e, PnC, CeM) within the *modulatory startle pathway-*the main focus of this work.

### Identification of the modulatory startle pathway

To investigate the neuro-functional basis of the *modulatory startle pathway*, we utilize two well-established experimental approaches for affect induction as an opportunity to obtain converging evidence for a common neural pathway of affect-modulated defensive responding in humans: The *affective startle modulation*(**ASM**) paradigm and a fear conditioning paradigm allowing for the investigation of *fear-potentiated* startle (**FPS**).

On a subjective and physiological level successful affect modulation was observed in both paradigms: In ASM, post-experimental *valence ratings* varied significantly for the three emotional picture categories [negative, neutral, positive; F(2,84)=398.88, p<0.001, η^2^=0.905, **Figure 2C**] in the expected directions (one-sided: negative<neutral, negative<positive, neutral<positive, all p<0.001). Accordingly, and replicating previous research outside the MR environment, *startle* eye-blink responses acquired during fMRI closely mirrored subjective valence ratings - commonly referred to as ‘affective startle modulation’ [F(2,68)=6.29, p=0.003, η^2^=0.156, **Figure 2C**]. More precisely, blink magnitudes were relatively potentiated during negative (one-sided: negative>neutral: p<0.043; negative>positive: p=0.001) and inhibited during positive picture viewing (one-sided: positive<neutral: p<0.030), hence following a valence-specific gradient of startle potentiation. In contrast, SCRs to picture onsets closely mirrored subjective *arousal ratings.* More precisely, significant differences across emotional categories [*arousal ratings*: F(2,84)=163.74, p<0.001, η^2^=0.796; *SCRs:* F(2,38)=6.31, p=0.004, η^2^=0.223, **Figure 2C**] reflect higher SCRs to emotionally salient (i.e., negative and positive) as compared to neutral pictures (*arousal ratings*: one-sided: negative>neutral, positive>neutral, both p<0.001; two-sided: negative vs. positive, p=0.200; *SCRs:* one-sided: negative>neutral: p<0.001, positive>neutral: p=0.003; two-sided: negative vs. neutral: p=0.541).

**Figure 2.**
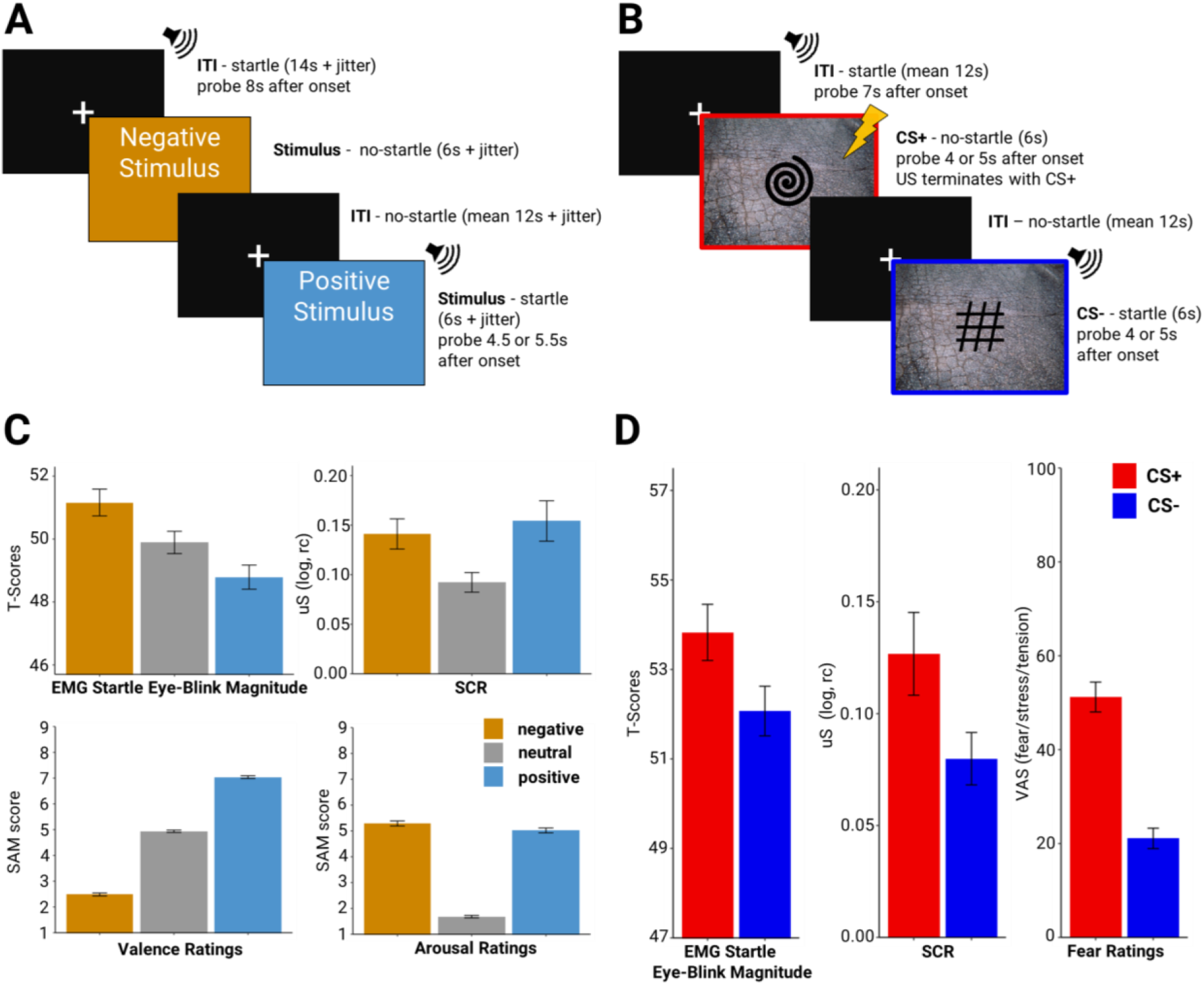
Experimental details of the employed paradigms. (A) Example of trial presentation for the affective startle modulation paradigm (neutral condition not shown). (B) Example trial presentations for the fear conditioning paradigm during the fear acquisition training phase (note that red and blue frames around the CS pictures serve illustrative purposes only). (C) ASM: mean responses during fMRI of startle eye-blink magnitude (affective modulation), SCR as well as post-experimental ratings. (D) FPS: mean responses during fMRI of startle eye-blink magnitude (fear-potentiated startle), SCR and subjective ratings of fear/stress/tension for CS+ and CS-in fear acquisition training. Error bars represent standard errors of the means.

In FPS, successful fear acquisition was indicated by significantly higher responses to the CS+ relative to CS-across all outcome measures: *fear ratings* [t(54) = 9.55, p<0.001], *startle eye-blink* [t(50) = 2.32, p = 0.012] as well as *SCRs* [t(43) =3.62, p < 0.001; **Figure 2D**].

In line with the observed valence-specific responding in subjective and psychophysiological measures, we observed stronger neural activation in PnC and CeM evoked by startle probes presented during unpleasant (ASM: negative>positive) and threatening (FPS: CS+>CS-) conditions, which are associated with potentiated startle eye-blink responses (**Figure 3A-D, Table 1**). Of note, mirroring the valence-gradient evident from both, startle eye-blink, valence and fear ratings, PnC and CeM activation followed the same pattern (**Figure 3EF**). In both studies, amygdala activation to the startle-eliciting stimulus seems to be markedly restricted to the central nuclei (**Figure 3CD)**, – the core output area of defensive responding and proposed key effector region of the PnC.

**Table 1.**
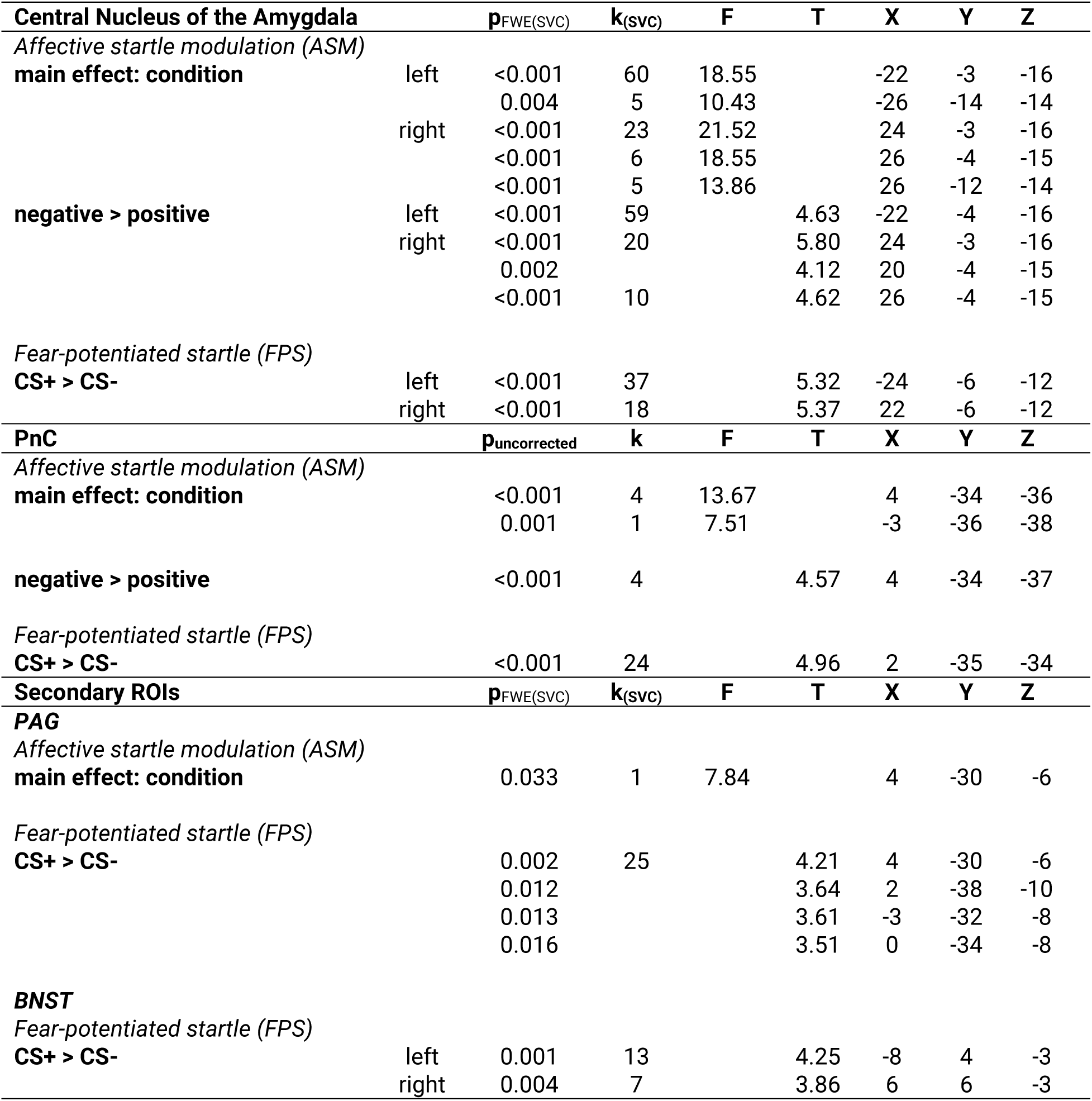
Statistics for Valence-dependent neural activation evoked by startle probes for ASM (F-Test main effect: condition, t-contrasts for negative > positive condition) and FPS (CS+>CS-) in both *apriori* defined regions of interest (CeM, PnC) as well as secondary regions of interest (PAG, BNST).

**Figure 3.**
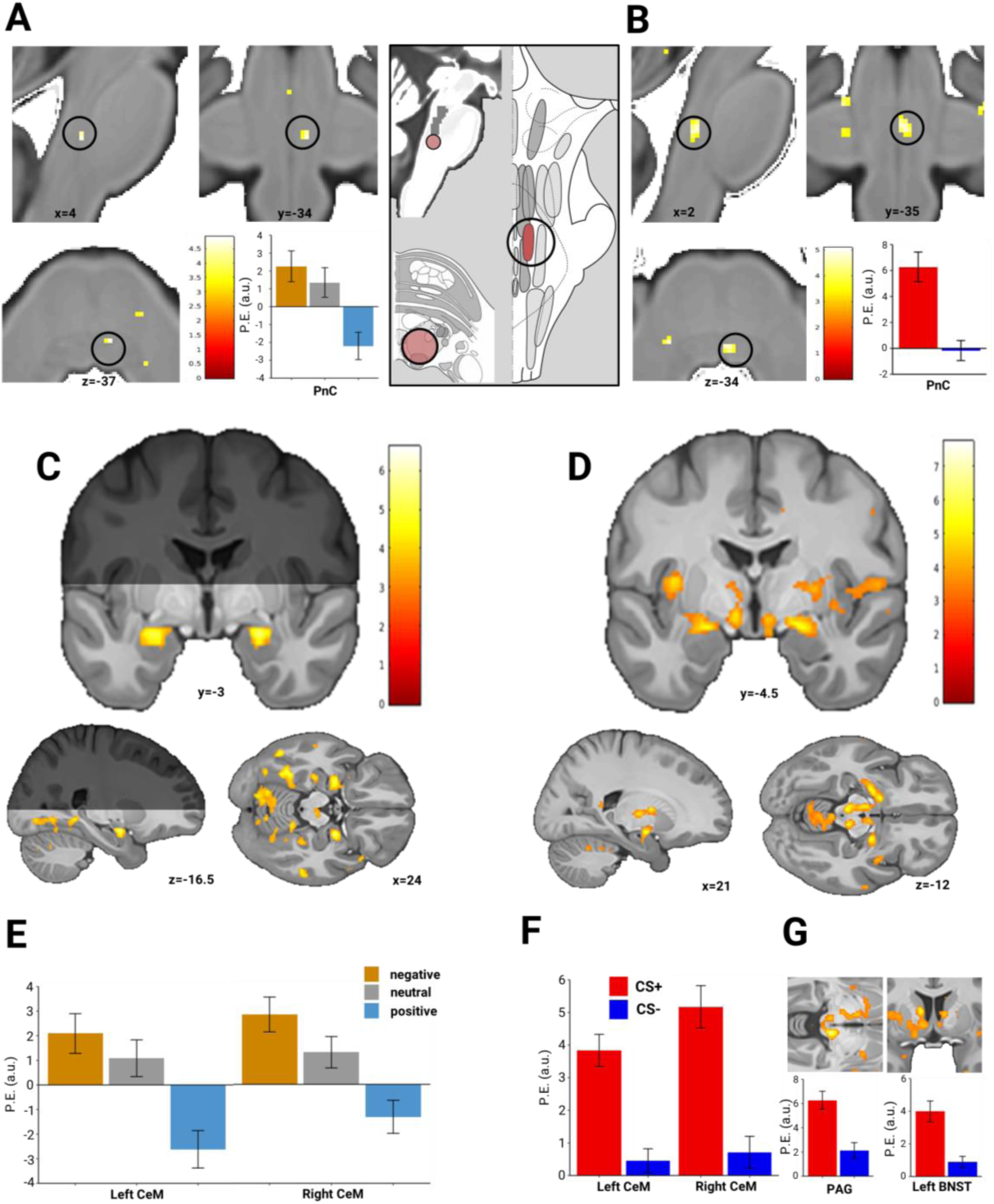
(A) Valence-dependent neural activation in PnC area and corresponding peak voxel parameter estimates evoked by startle probes during ASM: negative>positive (Note that black circles highlight activation within the PnC region and does not represent a specific size of search volume). (B) Valence-dependent neural activation evoked by startle probes in PnC area during FPS: CS+>CS- and corresponding parameter estimates of peak voxel results. Note that the grey box containing the schematic illustration of PnC area within Duvernoy’s Atlas of the Human Brain Stem and Cerebellum (Naidich et al., 2009, modified with permission) and the available anatomically defined MRI ROI of the PnO (Edlow et al., 2012) serves to illustrate overlap between expected area of the PnC and observed statistical maps. (C) Valence-dependent neural activation in the bilateral amygdala evoked by startle probes during ASM: negative>positive (the grey area illustrates restricted fMRI field of view) and (D) FPS: CS+>CS-.. (E) Corresponding parameter estimates extracted from peak voxel results in bilateral amygdala in ASM as well as (F) in FPS. (G) Valence-dependent neural activation of the PAG and the BNST in FPS and corresponding parameter estimates extracted from peak voxel. Display threshold at p_uc_<0.001. P.E. (a.u.): parameter estimates (arbitrary units). Error bars represent standard errors of the means.

In addition, in FPS, the BNST and PAG as our secondary ROIs were significantly implicated in fear-potentiated startle modulation (**Figure 3G, Table 1**). In ASM however, no valence-specific PAG activation was observed and the BSNT was not covered by the FOV (for details see **Table 1**).

In sum, we provide converging evidence for corresponding neural pathways underlying affect-modulated startle in rodents and humans – centering on the PnC and the CeM as key hubs.

### Trial-by-trial brain-behavior link during affect modulation of the startle reflex

An important further qualification of the observed valence-dependent responding on a psychophysiological and neural level can be established by *integrating* individual trial-by-trial EMG magnitudes into imaging analyses. These analyses quantify the linear relationship between EMG response magnitude and neural activation strength (i.e., parametric modulation) disregarding the categorical valence information within the statistical model.

Acquisition parameters and the design of ASM was specifically tailored to enable these methodologically inherently challenging analyses while this question can only be addressed in an exploratory way in FPS.

In ASM, trial-by-trial magnitudes were indeed reflected in activation strength of the PnC (p_uc_ = 0.001, T = 3.65, k = 1, [x,y,z] = [2,-35,-34], **Figure 4A**) as well as left CeM (p_SVCFWE_ < 0.001, T = 3.76, k = 5, [x,y,z] = [-24,-1.5,-16.5], **Figure 4A;** p_SVCFWE_ = 0.001, T = 3.56, k = 1, [x,y,z] = [-18,-6,-14]) providing a hitherto direct link between defensive behavior and corresponding neural activation.

**Figure 4.**
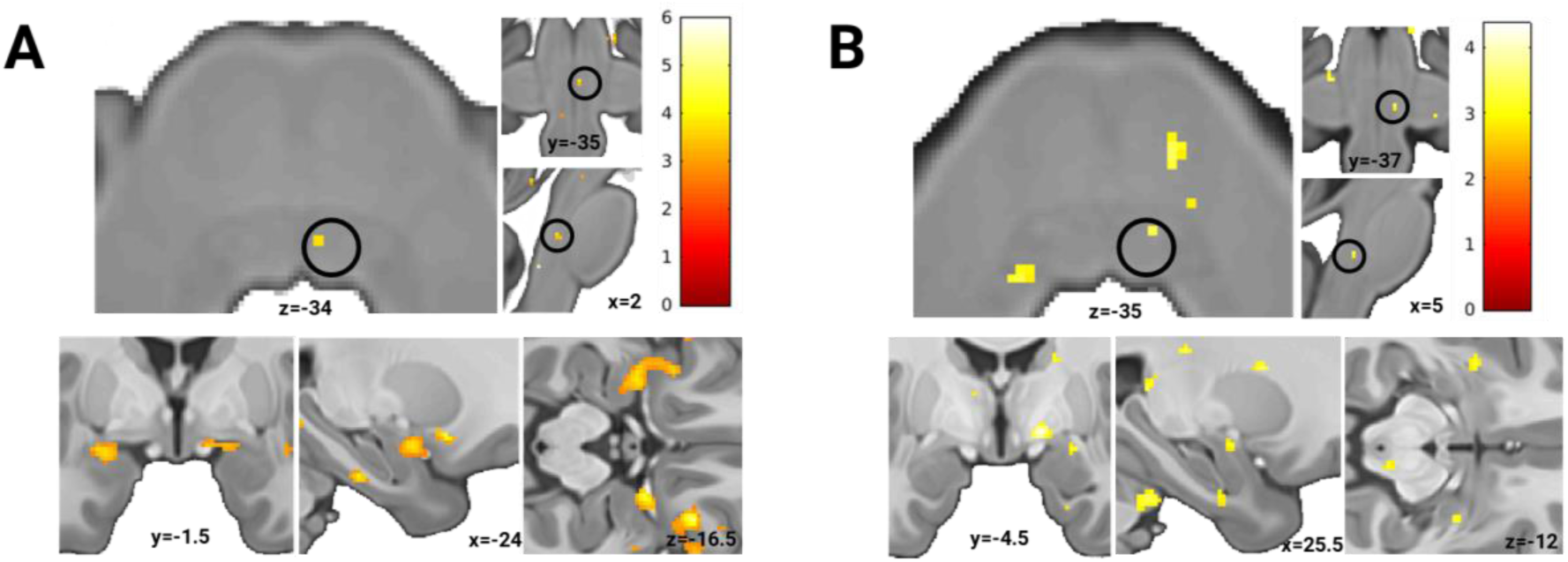
Activation of the PnC and CeM functionally mirroring trial-by-trial EMG magnitudes per individual in (A) ASM and (B) FPS. Display threshold at p_uc_<0.005.

**Figure 5.**
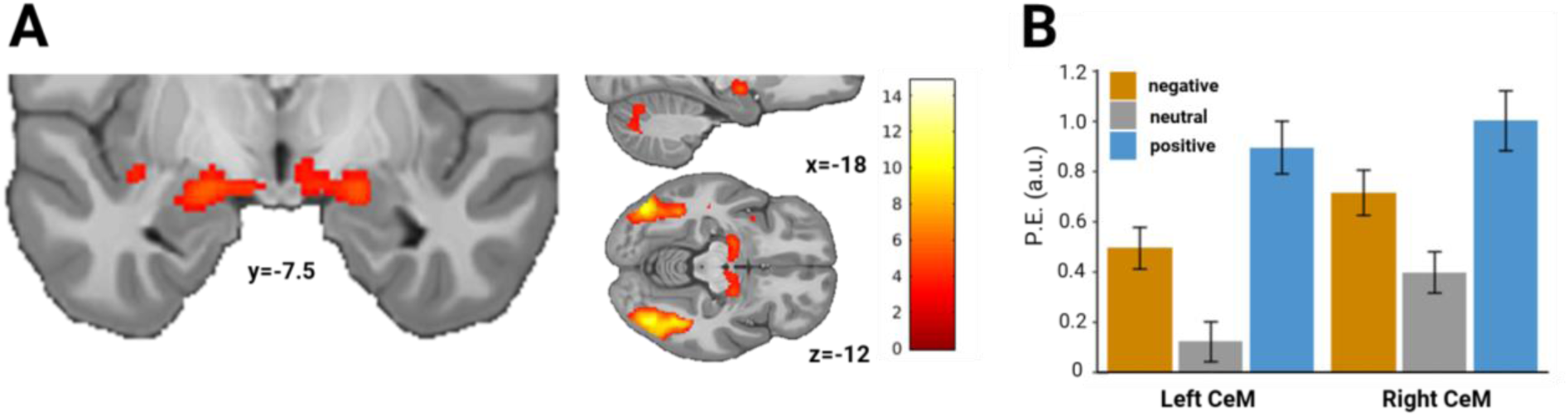
(A) Arousal-like pattern in the amygdala during picture viewing (emotional pictures>neutral) in the ASM study. Note that, for these analyses, only trials without startle probes were used to avoid confounding of picture viewing-related activation by activation related to startle probe presentation. (B) Extracted parameter estimates in left and right CeM. Display threshold at p_uc_<0.001. P.E. (a.u.): parameter estimates (arbitrary units). Error bars represent standard errors of the means.

Exploratory analyses of FPS support this association within the PnC (p_uc_ = 0.003, T = 3.20, k = 2, [x,y,z] = [5,-35,-37], **Figure 4B**) and the right CeM (p_SVCFWE_ = 0.002, T = 3.01, k = 2, [x,y,z] = [25.5,-4.5,-12], **Figure 4B**) albeit at a more liberal threshold of p<0.005uc. Importantly, correspondence of significant peak voxel coordinates across both studies and analyses approaches (i.e. pre-defined categorical affective conditions vs. parametric EMG data integration) increases confidence in the observed relationship in the PnC and CeM.

### Dissociation in amygdala activation during passive and triggered responding

Our observation of *valence*-specific *triggered* CeM responding (i.e., evoked by the startle-eliciting stimulus, **Figure 3CE**) is intriguing, since it stands in marked contrast to the commonly observed *arousal*-dependent amygdala responding ^37^during *passive* emotional picture viewing.

Importantly, investigating *passive* processing (i.e. passive viewing) of emotional pictures in our data replicates these previous reports of an arousal-dependent response pattern in SCR, arousal ratings (see **Figure 2B**), and importantly also bilateral CeM activation (**Figure 5AB**, F-test; left: p_SVCFWE_ < 0.001, F = 21.12, k = 55, [x,y,z] = [-18,-7.5,-12]; right: p_SVCFWE_< 0.001, F = 17.07, k = 9, [x,y,z] = [19.5,-9,-13.5] and p_SVCFWE_= 0.003, F = 10.45, k = 3, [x,y,z] = [19.5,-4.5,-15], **Figure 5**).

In sum, we observe a marked dissociation between CeM responding to passive processing of emotional information which follows an arousal-like pattern (negative and positive > neutral) and triggered CeM responding elicited by startle probes presented on emotional foreground information which follows a valence specific pattern (negative > neutral > positive).

## Discussion

Here, we utilized recent advances of combined EMG-fMRI and brainstem imaging to delineate the neural pathway underlying the affective modulation of the acoustic startle reflex in humans and provide the critical direct brain-behavior link across two independent samples and experimental paradigms (i.e., affective startle modulation, ASM, fear potentiated startle, FPS). In agreement with rodent work, we provide converging evidence for a conserved underlying neural pathway in humans centering on the PnC and the CeM. Our results further highlight the value of combining startle eye-blink EMG with fMRI measurements as a unique opportunity to probe valence-specific triggered amygdala responding as a promising novel read-out measure in affective neuroscience.

The PnC functions as key hub in the primary acoustic startle reflex ^5,6^for initiating the startle response and for integrating affective information. Here, we demonstrate startle-evoked neural responses in the PnC also in humans. Most importantly, we show that PnC activation is indeed modulated by affective input, presumably transmitted from the CeM. On a defensive response level, this manifests as affective modulation of the startle eye-blink EMG response magnitude. In addition to these key findings, we show a startle-evoked affective modulation of BNST and PAG activation in the FPS study that involved imminent threat. This corroborates their proposed involvement in the processing of fear-related information ^5^and substantiates their role in defensive responding (i.e., protective reflexes such as startle), which may motivate further detailed investigations.

An important qualification of the identified affective modulation of PnC and CeM activation is the demonstration of a direct trial-by-trial brain-behavior link relating strength of neural activation to individual EMG eye-blink startle magnitudes in these key hubs of the modulating pathway. As current evidence for an association between affective (i.e., fear) modulation of the startle response has been based on lesion studies only ^28–30^, these findings provide an important direct and novel link quantifying the relationship between eye-blink response magnitude and neural activation strength in the brainstem (i.e., PnC) as well as the centromedial part of the amygdala (CeM).

Critically, our results demonstrate an important dissociation between neural mechanisms of cue-related emotional processing and the startle reflex itself: In the behavioral lab, the dissociation between eye-blink EMG response and skin conductance responses, which mirror valence-specific and arousal-specific responding respectively ^13,38^is well described. Importantly, combining eye-blink EMG with fMRI acquisition now allowed us to demonstrate this dissociation at a neural level. In detail, we observe the expected arousal-specific CeM responding (i.e. emotional > neutral) during emotional picture viewing ^37^in the ASM study, which closely follows skin conductance responses and is in line with a role of the amygdala of allocating attention to salient signals ^39–42^. Importantly, however, this response pattern in the CeM switches to a valence-specific responding, mirroring startle responding, through presentation of the auditory startle probe – an external event triggering defensive behavior. More precisely, depending on the affective state induced by the picture itself, CeM activation triggered by the startle-eliciting stimulus was either potentiated when presented on negative background information or inhibited when presented on positive background information. This observation crucially supports the proposed function of the amygdala as gatekeeper for coordinated responses after initial evaluation of stimulus threat value ^1^. This pattern of observation is both intriguing and potentially highly relevant for future work on valence-dependent processing ^43^. This triggered amygdala output can be expected to mirror (observable) defensive responses towards potential threat more closely than measuring tonic amygdala responding elicited by emotional processing. Hence, such triggered events may function as a read-out of the ‘state’ of the amygdala, which might not be accessible otherwise. As such, we suggest that *triggered* amygdala responding may prove as a useful tool in the future.

In line with this, our results highlight the value of combining startle eye-blink EMG with fMRI measurements to provide a new ^22–26^and, importantly, valence-specific read-out measure for affective and clinical neuroscience. Hitherto, studies have primarily used SCRs or pupil dilation in the MRI context, which however, capture arousal but not valence-specific gradients ^13^. In particular, the observed direct relationship between startle eye-blink EMG magnitude and neural activation strength on an individual level presents a potential opportunity to use individual startle eye-blink measures as direct read-out of neural activation of the central amygdala and the brainstem nuclei priming the motor output. In particular, in light of recent calls for investigations of responding beyond the average this is in line with the current zeitgeist ^44^to enable investigations on an individual trial-by trial level. As, more precisely, a limited focus on average responding may, at best, deprive us from crucial insights into the underlying mechanisms as average responding may in fact represent an artificial pattern that does not reflect the response of any individual subject well (for instance in the case of unrecognized subpopulations).

Some limitations of our work are, however, worth noting: First, defining the exact anatomical location of most brainstem nuclei is a challenge as anatomically defined boundaries are not available. Yet, we provide both spatially and functionally converging evidence from two independent samples and paradigms that support the accuracy of the PnC location in our work. Hence, we provide anatomical coordinates that future work may utilize for defining the PnC area.

Second, in the FPS study, the startle probe regressor in the fMRI model shows inherent collinearity with the US regressor due to its close proximity in time. While the jittered startle probe onset as well as its presentation in only 66% of trials already reduces interpretation problems, this collinearity occurs only on the first (i.e., individual) level. Importantly, collinearity at the first level (as opposed to the second level) is not subject to estimation problems or increased risk of false positives but may result in highly variable parameter estimates and hence decreased sensitivity ^45^.

Third, our results are exclusively based on startle responding triggered by acoustic stimuli and it can hence only be speculated that our results also generalize to other trigger modalities (e.g., tactile, visual).

Fourth, we employed a strictly hypothesis-driven region-of-interest approach. Future studies can now build on these results in formulating and testing additional hypotheses also involving other regions of interest. For instance, our work, and hence the FOV covered in the ASM study, primarily focused on potentiation of the startle reflex. Inhibition of the startle reflex might be of interest as well but regions potentially involved in this processes were not assessed due to the acquisition area tailored to subcortical and brainstem ROIs in the ASM paradigm. In fact, an expression of the full startle response is energetically costly and has been suggested to lead to potential foraging opportunity losses ^46^. The new methodological avenues opened up by our work can now be used to explore this further.

In conclusion, in human affective neuroscience, reflexive responding and its adaptation to environmental demands has hitherto not received much attention (however, see ^47^) - in contrast to higher order (cognitive) components of emotional processing and regulation. By highlighting the cross-species conserved neural pathway of defensive startle reflex modulation, we provide an important yet missing piece connecting hitherto separate lines of research on 1) the role of the amygdala in emotion processing in humans (e.g. fear learning) and 2) the role of the amygdala in affective startle reflex modulation in rodents. This corroborates the role of startle reflex modulation as *the*prime cross-species translational tool of defensive reactivity in affective neuroscience ^48,49^. Critically however, its application in humans has been limited to behavioral work by technical and methodological constraints in the past. Here, we demonstrate both the applicability of EMG eye-blink startle responding in the fMRI context and provide the crucial direct brain-behavior link for affective startle modulation. This will allow to explore entirely new avenues in the future that can be expected to provide major novel insights in affective neuroscience.

## Acknowledgements

The authors thank Christian Möller, Jürgen Finsterbusch and Katja Hillbrandt for technical support for preparing EMG-fMRI data acquisition, Maike Möller and Jana Hofacker for help with participant preparation and data acquisition, Christian Sprenger for help with data analysis, Katrin Bergholz and Kathrin Wendt for technical assistance during MR data acquisition, as well as Jan Haaker and Christoph Korn for comments on previous versions of this manuscript.

The author thank the University Medical Center Hamburg Eppendorf (Forschungsfond Medizin), the Deutsche Forschungsgemeinschaft (DFG, German Research Foundation) – Projectnumber 44541416 – TRR 58 (sub-project B07) to TBL.

## Author contributions

M.K., J.W., A.H., and T.B.L conceived and designed the study. M.K., J.W., R.S. and T.B.L. designed and implemented the experiments. M.K., and R.S. acquired data. M.K., C.B., and T.B.L. analyzed data. All authors have contributed to results interpretation and drafting of the manuscript.

## Methods

### Subjects and experimental design

All subjects were recruited via online advertisement and provided written informed consent. Protocols were approved by the ethics commission of the German Psychological Society (DGPs) (**ASM**) or the General Medical Council Hamburg (**FPS**).

#### Affectivestartlemodulation (ASM)

Forty-three male subjects [mean age (s.e.): 25.88 (0.41)] were recruited. A standard affective pictures startle modification task^9^ was employed (**Figure 2A**). Standardized and validated (for arousal and valence) pictures from the International Affective Pictures System (IAPS,) as well as from the EmoPicS database^50^ were used. 12 pictures per category [negative (human attack), neutral (depicting humans), positive (erotica of couples and single opposite sex)] were selected based on matched valence and arousal ratings per category to robustly elicit a reliable affective startle modulation (i.e. inhibition and potentiation). Only male subjects were included to avoid sex-specific stimulus selections since erotic pictures served as the positive stimulus category. Pictures with comparable social content were selected. The paradigm was validated in a preceding behavioral pilot study in an independent sample (N_pilot_= 24, data not shown). Pictures were presented for 6s separated by randomized inter-trial-intervals [ITIs, durations 10, 12 or 14s]. Moreover, a jitter (0, 1/4, 1/2 or 3/4 of a TR) added to the ITIs allowed for oversampling of the hemodynamic response function (HRF) of the BOLD-signal. ITI and jitter length were counterbalanced between trial lists and picture valence. Picture valence was counterbalanced across trial lists. Each category was shown maximally for two consecutive presentations and transition likelihood between categories was kept constant. Each picture was presented twice and startle probes were presented in 50 percent of picture presentations at 4.5s or 5.5s (counterbalanced across categories and trial lists) after stimulus onset. To avoid predictability of the startle probes, 12 startle probes were added across ITIs each occurring during one of the 14s ITI (+added jitter) periods with onset 8s after ITI onset. During an initial startle habituation phase (**Figure 1B**), eight startle probes were presented while displaying a fixation cross as shown during the ITI. Habituation startle probes were separated by 11s (+ added jitter). The separation of the habituation startle probes by this long inter-stimulus-interval (ISI) and the addition of the jitter particularly allowed to quantify the individual neural response to each startle probe (which was not possible for the short ISIs in the FPS startle probe habituation phase).

To increase subjects’ alertness, an ‘oddball task’ was included. Subjects were instructed to press a button whenever a scrambled picture was presented. These pictures were taken from the neutral picture group, scrambled in cubes (25-by-25 pixels in size) and not recognizable in content.

In total, each picture per category was presented twice with three additional oddball presentations resulting in 75 trials.

Pictures (800×600 pixels) presented on a grey background were projected onto a screen (1024×786 pixels) at the back of the magnet’s bore within the MR scanner which participants could see via a mirror mounted over their heads. Visual and auditory stimuli were presented using Psychophysics Toolbox-3 ^51^running on MATLAB2010b (The MathWorks, Natick, MA, USA).

Post-experimental ratings for valence and arousal using the self-assessment manikin scale (SAM, ^52^) were employed (see below for details).

#### Fear-potentiatedstartle (FPS)

Fifty-five subjects [female = 36; mean age (s.e.): 25.6 (0.47)] were recruited. The fear conditioning paradigm (experimentally similar to Sjouwerman et al. (2016)^53^, **Figure 2B**) consisted of six phases: startle probe habituation to achieve a stable baseline for startle reactivity, CS habituation, fear acquisition training, immediate extinction, reinstatement, and a reinstatement test. The present study focuses on the startle probe habituation, CS habituation and fear acquisition training phases only and hence we provide no further details with regard to the other experimental phases. Prior to the experiment, participants were explicitly instructed to not attend to the startle probes to avoid interference with CS-US contingency acquisition. No explicit instructions regarding the CS-US contingencies were provided.

During startle probe habituation, five startle probes (ISI: 6s) were presented while displaying a white fixation cross on black background which was also shown during the inter-trial interval (ITI, durations 10, 11, 12 or 13s) during the following experimental phases. Two geometric shapes (spiral and hash) served as CS stimuli and were displayed on a background image (water, sand, grass or concrete). During CS stimuli habituation, both CSs were presented twice. During fear acquisition training, CSs were presented 9 times each. One of the CSs (CS+) co-terminated with an electro-tactile US (100% reinforcement rate), whereas the other CS was never paired with an US (CS-). Allocation of the CS to the CS+ and CS- and the order in which the CS+/CS- appeared were counterbalanced between individuals. The startle probe was delivered for half of the CS stimuli during CS habituation (i.e., one for CS+, one for CS), for two thirds of the CS stimuli during fear acquisition training (4 or 5s after CS onset), and for one third of all ITIs (5 or 7s after ITI onset). The first and last CS presentations per trial type during fear acquisition training were always presented with a startle probe. Presentation of all stimuli was controlled using Presentation Software (NeuroBehavioral Systems, Albany, CA). Visual stimuli were projected onto a screen (1024×786 pixels) at the back of the magnet’s bore within the MR scanner which participants could see via a mirror mounted over their heads.

The unconditioned stimulus (US) was administered as an electro-tactile stimulus consisting of a train of three 2-ms square waves with an ISI of 50ms to the back of the right hand. The electrical stimulation was generated by a DS7A electrical stimulator (Digitimer, Welwyn Garden City, UK) and delivered through an electrode with a platinum pin surface (Specialty Developments, Bexley, UK). Prior to the experiment, US intensity was calibrated for each participant individually [unpleasant but tolerable, aiming at an intensity of 7 out of 10 (with 10 referring to the most unpleasant sensation that might be inducted by the electrode), mean intensity (s.e.) of the final sample (N=55): 5.12mA (0.47)]. Intermittent ratings of fear/stress/tension were acquired (see below for details).

### Subjective ratings

#### ASM

Outside the MR-environment subjects rated each picture at a computer screen (screen size 1920×1200 pixels, stimulus size 800×600 pixels) after scanning. A 9-Point Self-Assessment-Manikin (SAM,^52^ rating scale for valence (from 1 = very pleasant to 9 = very unpleasant) and arousal (from 1 = very calm to 9 = very arousing) was used. After one initial training trial using a novel neutral stimulus to familiarize the subject with the rating procedure, all pictures from all three categories were consecutively presented at random with rating scales of valence and arousal, respectively, beneath the picture. Ratings were selected via mouse click at the subject’s own pace and rating times were recorded to check for compliance of the subject.

#### FPS

Participants indicated their level of fear, anxiety, and distress toward both CS types (“How much stress, fear, or anxiety did you experience the last time you saw symbol X?” with the X referring to one of the CS types at a time) intermittently throughout the experiment (in one rating block after CS habituation and three rating blocks during fear acquisition training) on a visual analogue scale (VAS) ranging from 0 (none) to 100 (maximum). A rating block was preceded by a screen that signaled the start of the rating block for 4s. The rating block included one rating for the CS+ and one for the CS- in a randomized order. The start position of the curser was randomly placed on the VAS for every trial. Participants were required to confirm their rating within 9s otherwise the rating trial was regarded as invalid and treated as missing. Ratings within blocks were separated by an ITI of 1s. An additional rating for the aversiveness of the startle sound was included after the habituation phase.

### Psychophysiological data acquisition and processing

For both studies, electromyography (EMG) startle eye-blink and skin conductance responses (SCR) were acquired. Data acquisition and processing were identical across studies.

#### EMG

In both studies, startle eye-blink response data were acquired through EMG recordings within the MR environment using a FaceEMG Cap-MR (EasyCap GmbH, Herrsching, Germany). The cap contains five Ag/AgCl electrodes and built-in 5 kOhm resistors to facilitate wire management in the magnetic field, to ensure participant and equipment safety as well as artifact reduction in EMG and BOLD data. Skin was prepared with abrasive electrode gel. Two electrodes were placed at the participant’s orbicularis oculii muscle beneath the left eye; two electrocardiogram (ECG) electrodes were placed at the participant’s back and one ground electrode was attached to the forehead of the participant. The maximum transmission resistance threshold was set to 20 kOhm.

A 50ms burst of white noise served as startle probe which was calibrated to 103dB[A] using a MR-compatible sound level meter (Optoacoustics Ltd., Mazor, Israel) and presented to subjects binaurally via headphones (MR Confon GmbH, Magdeburg, Germany). Baseline scanner noise during EPI acquisition in both studies was approximately 79dB[A]. EMG data were recorded within the BrainVision Recorder software using BrainAmp ExG MR amplifier (Brain Products GmbH, Gilching, Germany), including the SyncBox device for synchronization of recorded data and MR gradient switching ^54^, applying a 16 bit Analog-to-Digital-Conversion (ADC). Sampling rate was set to 5 kHz with a signal resolution of 0.1 μV within a frequency band of 0.016 and 250 Hz.

Data were processed as described previously ^24^. Briefly, this included MR gradient correction by subtraction of an artifact template averaged over seven EPI volumes from the raw signal, down-sampling to 1000 Hz, reduction of eye-movement related artifacts by application of a low cutoff filter of 60 Hz with a time constant of 0.0027 and 48 db/oct, rectification, and manual offline scoring using a custom-made computer program. Following published guidelines ^8^, the magnitude of the eye-blink response (in microvolts) was measured from onset to peak, as described previously ^55^. Eye-blink magnitudes were T-transformed (including all experimental phases and conditions, see below) for statistical analyses of startle responses while raw values were fed into fMRI trial-by-trial first-level analyses (see fMRI analyses for details). Undetectable blinks were scored as zero responses and as missing if a blink occurred immediately (up to 50ms) before startle probe administration or due to excessive baseline activity, obvious electrode, or gradient artefacts.

#### SCR

In both studies, SCRs were measured via self-adhesive Ag/AgCl electrodes which were placed on the palmar side of the left hand on the distal and proximal hypothenar. Hands were washed with tap water and without soap. Data were recorded with a CED2502-SA skin conductance unit together with a Biopac MP150-amplifier system (BIOPAC Systems Inc, Goleta, California, USA) with Spike 2 software (Cambridge Electronic Design, Cambridge, UK). Data were down-sampled to 10 Hz, smoothed by using a 5-point moving average and phasic SCR to stimulus onsets were manually scored offline using a custom-made computer program. SCR amplitudes (in µS) were scored as the largest response initiating 0.9 to 4.0 s after stimulus onset ^56^. Non-responses were scored as zero and trials showing recording artefacts were scored as missing data. Logarithms were computed for all values to normalize the distribution ^57^, and these log values were range-corrected (SCR/SCRmax) to account for inter-individual variability ^58^.

#### SubjectpreparationandEMGdataqualitycontrol

In both studies in the scanner, SCR electrodes and a respiration belt were attached, a pulse-oximeter was attached to the left index finger and headphones were placed on the subject’s head. Afterwards, the EMG cap was connected to the EMG amplifier. We ensured no heat build-up in the electrodes and that the subject’s visual area was not restricted by the EMG equipment placed behind the projection screen. Subsequently, impedances of the EMG/ECG-electrodes were re-checked to ensure subject’s safety. SCR, pulse and respiration signals were visually inspected to check for data quality of physiological responses prior to the experiment. The auditory startle probe was presented to ensure the subject’s compliance with the sound level of the stimulus and to check data quality of the EMG signal without gradient artifacts of the scanner (i.e. scanner offline). Following this set-up procedure, a structural image (see MRI acquisition) was acquired to allow the subject to get used to the environment and to ensure all equipment was working safely. The subject was reminded that from time to time an auditory stimulus will be presented without any relevance to the stimuli. The subjects was informed that the experiment will begin with the presentation of several (ASM: eight / FPS: five) of the auditory stimuli before the first visual stimuli are presented (i.e. startle probe habituation phase). Starting with the scanner gradient, the EMG signal was visually inspected via online MR-artifact correction with Brain Vision’s RecView to ensure all electrodes were still attached and signal quality was good.

### Data analyses of ratings and psychophysiology

Data analyses were homogenized across both studies whenever feasible. Insufficient data quality (defined by more than 66% of missing values or null responses for ASM: during startle habituation phase and the actual experiment; for FPS: during startle habituation, CS habituation and fear acquisition training) led to exclusion of data from eight subjects for EMG analyses and 23 subjects for SCR analyses in the ASM study and four subjects for EMG analyses and data of eleven subjects for SCR analyses in the FPS study.

#### ASM

Repeated-measures analyses of variance (rmANOVA) were performed in R ^59^to assess the differences between categories (negative, neutral, and positive) as within-subject factor for ratings of valence and arousal as well as EMG responses and SCRs (effect sizes reported as partial η^2^). Significant effects were followed up via post-hoc t-tests to specify differences across categories. While for valence ratings and EMG responses one-sided t-tests were performed as strong a priori assumptions exist for the direction of effect (i.e. positive>neutral>negative for valence ratings; negative>neutral>positive for EMG responses), for ratings of arousal as well as SCRs, one-sided t-tests were only performed for comparisons between emotional and neutral conditions whereas a two-sided t-test was performed between both emotional categories because we had no hypothesis regarding differences between the emotional categories (negative vs. positive).

#### FPS

For ratings, EMG and SCR measures, one-sided paired-sample t-tests were performed in R ^59^to investigate the differences of mean responding towards CS+ and CS-during the fear acquisition training phase.

### Functional magnetic resonance Imaging (fMRI)

#### Dataacquisitionandprocessing

For both studies, MR data were acquired on a 3T MRI scanner (MAGNETOM Trio, Siemens, Erlangen, Germany) using a 12-channel head coil. A high-resolution T1-weighted structural image (1×1×1mm) was acquired using a magnetization prepared rapid gradient echo sequence (MPRAGE).

#### ASM

Imaging parameters were specifically tailored to the brainstem and amygdala as our prime regions of interest: Twenty-five continuous axial slices (2 mm thick, no gap) were acquired using a T2*-sensitive gradient echo-planar imaging (EPI) sequence [repetition time (TR): 2.0 s; echo time (TE): 27 ms; flip angle: 70°; field of view (FOV): 232 x 232 mm, 2 x 2 mm in-plane resolution] in three sessions. TE was minimized using a parallel acquisition technique (generalized auto-calibrating partially parallel acquisitions, GRAPPA) with an acceleration factor of 2 and 24 reference lines. The slice package was adjusted to cover the lower border of the pons and the amygdala on the upper side. To avoid scanner drift artifacts over time, the experiment was divided into three scanning sessions in between which the scanner was re-adjusted without any interaction with the subject.

#### FPS

Imaging parameters were selected to cover the brainstem and simultaneously achieve near-complete coverage of the brain. 37 continuous axial slices (2 mm thick, 1mm gap) were acquired using a T2*-sensitive gradient echo-planar imaging (EPI) sequence [TR: 3.0 s; TE: 26 ms; flip angle: 90°; FOV: 220 x 220 mm, 2 x 2 mm in-plane resolution]. TE was minimized using GRAPPA with an acceleration factor of 2 and 48 reference lines.

Data was processed within SPM12 (http://www.fil.ion.ucl.ac.uk/spm/) running on MATLAB2013a (The MathWorks, Natick, MA, USA). Initial fMRI preprocessing steps included discarding the first four volumes of each time series to account for T1 equilibrium effects, slice-time correction [for ASM (Study 1), using a HRF oversampling protocol], realignment and motion correction using the unwarp function implemented in SPM12, manual inspection for excessive head movement and co-registration with the structural image. Two spatial normalization procedures of functional data were employed to account for standardized cortical imaging and the specific requirements for brainstem imaging, respectively. First, for analyses of the whole coverage of the FOV, data was normalized using DARTEL ^60^and spatially smoothed with an isotropic Gaussian kernel (6mm FWHM) for analyses of whole brain effects with specific focus on the amygdala as region of interest). Second, for improved brainstem spatial normalization, data was normalized using the SUIT toolbox as implemented for SPM ^61^including up-sampling the data to a resolution of 1×1×1m. Since the target brainstem region of interest (i.e. the nucleus pontis caudalis, PnC) is a very small nucleus located within the pons, no spatial smoothing of functional data was applied during brainstem specific normalization ^34^. Note that all presented coordinates obtained from the brainstem specific analyses are in reference to the space defined by the SUIT toolbox and are in close alignment with the MNI space. During statistical estimation, further processing included temporal high-pass filtering (cut-off 128s) and correction for temporal auto-correlations using first-order autoregressive (AR1) modeling. Additionally, for brainstem specific statistical analyses, physiological noise correction was performed by adding 18 regressors of no interest using RETROICOR ^62^which were estimated based on individual physiological data of cardiac (pulse curve recorded via pulsoxymeter) and respiratory data both acquired with a MR compatible monitoring system (Expression, InVivo, Gainesville, USA).

For illustrative purposes, parameter estimates of analyses were extracted using the rfxplot toolbox as implemented in SPM ^63^.

#### Dataanalysesofprimarystartlepathway

The neural response to *startle probes* was investigated to explore the involvement of the PnC in the primary pathway prior to investigating the modulatory pathway of the startle reflex, which was the focus of this work.

The startle habituation phase of the ASM paradigm was particularly suited to investigate neural responding towards repetitive startle probe presentation (**Figure 1B)** to identify PNC involvement in the primary pathway because (1) no meaningful visual stimuli are presented during this phase and (2) the timing of presentations (i.e. 11s between startle probes plus jitter of 0, 1/4, 1/2 or 3/4 of a TR) allowed to separate the neural responses to these probes. Hence, this analysis is based on the ASM paradigm only. This analysis is based on eight habituation trials included in the first-level models described below for the valence-specific categorical analyses. Thereby, the parameter estimated for the onset reactivity for all eight probes is taken from the first-level to a one-sample t-test for second-level statistics. A directional contrast testing for positively associated activation with probe onsets was used to assess the neural responding towards the habituation startle probes. In addition, to explore a direct brain-behavior link, EMG data were combined with the fMRI data (i.e. parametric modulation). Therefore the first-level model was extended by modulation of the habituation phase startle probe regressor with values of the time-dependent EMG habituation pattern that was observed across all participants (**Figure S1B**). In this approach trial-by-trial mean T-transformed EMG amplitudes were used in order to compensate for the limited number of trials and missing data. Note, this pattern is based on a mean responding of participants and does not include individual responses, as this analysis is based on only eight data points and missing data would hence reduce sample size and sensitivity within subjects. Estimated parameters for the modulated regressor are taken to the second-level by means of a one-sample t-test investigating a potential link between the neural activation within the regions of interest and the time-dependent EMG response pattern.

#### Data analyses of modulatory startle pathway

For both studies, a two-step approach to analyzing *neural responses to startle probes* was employed using 1) valence-specific categorical and 2) EMG signal-integrative (i.e., parametric) analyses. First, the valence-specific categorical approach comprises *average* (i.e., across subjects) neural responses to startle probes for all affective conditions (including subjects with insufficient EMG data quality). Second, this was complemented by analyses directly linking neural activation to the individual EMG amplitudes. Here, preprocessed trial-by-trial eye-blink data *onanindividual*basis was integrated into an fMRI analyses as a parametric regressor.

#### Valence-specific categorical analyses

For the ASM paradigm, a general linear model (GLM) was set up for statistical first-level analysis including six regressors combining stimulus category and startle condition (i.e., no-startle-negative, no-startle-neutral, no-startle-positive, startle–negative, startle-neutral, startle-positive). Stimulus presentations were modeled as continuous blocks while overlayed startle probes were modeled as events. Two additional regressors for the habituation startle probes and inter-trial startle probes were modeled as events. Additionally, one block regressor for three oddball-trials (see SI for details) was added. To allow for the integration of the continuous EMG signal across sessions into of the fMRI data, data of all three sessions were concatenated. All regressors were convolved with a canonical HRF. Second-level analyses used SPM’s flexible factorial model which permits correction for possible non-sphericity of the error term and takes a subject factor into account when analyzing differences in within-subject conditions. Within this framework, first, a non-directional F-test was carried out to investigate differential neural activation to startle probes within the regions of interest (see below) between all three valence conditions (i.e., main effect: condition). Following up on these results, particular interest was directed at the *a priori* expected neural activation related to sub-cortical and brainstem responding to the startle probe during emotional conditions of negative-valence states (i.e. startle potentiation) as compared to positive-valence states (i.e. startle inhibition). Therefore, follow-up directional contrasts for neural responses towards the startle probe in negative>positive conditions were investigated. To explore the neural response to emotional pictures, an additional non-directional F-test based on a flexible factorial model, including estimated parameters for the emotional condition blocks, was calculated.

For the FPS paradigm, a GLM was set up for statistical first-level analysis. Regressors were constructed for CS onsets combining CS-type (CS+/CS-) as well as startle presentation (no-startle/startle) during the CS habituation as well as the fear acquisition training phase, respectively. For CS habituation, these regressors served as regressors of no interest. Moreover, four additional regressors modeling the onsets of the habituation startle probes, inter-trial startle probes during CS habituation, inter-trial startle probes during fear acquisition training as well as for the USs were built. Ratings across all phases were modeled in one regressor as blocks for the entire duration of each rating block. All regressors were convolved with a canonical HRF function. *Apriori* directional t-contrasts were calculated for the hypothesized effect of interest for increased startle probe onset reactivity during CS+ (threatening/stressful) as compared to CS-(safe/not stressful) conditions (i.e., CS+>CS-) within the fear acquisition training phase. Second-level analysis used SPM’s one-sample t-test to test for significant differences across all individuals within the pre-defined regions of interest (see below).

#### EMG signal-integrative parametric analyses

First-level models designed for integrated eye-blink response data were similar to both models used in categorical analyses for ASM and FPS. However, for both studies, onsets for all startle probe regressors contained in one design matrix (i.e., startle probe onsets during conditions, startle probe habituation, and inter-trial-intervals) were condensed into one single regressor of interest. To assess the correlative relationship between neural and muscular activation, recorded raw EMG magnitudes were used as parametric modulator of the startle probe onset regressors. Raw EMG magnitude values were used because of the summary statistics approach and the centering procedure implemented for parametric modulators within SPM12. When blink responses were classified as missing values, these startle probe onsets were excluded from the startle regressor of interest and added to the design model as single regressor of no interest. To guarantee a stable parameter estimation and thereby a meaningful association between EMG magnitude values and neural activity, subjects having more than 33% missing values within the entire experimental phase were excluded from further second-level analyses. This led to reduced sample sizes for second-level analyses based on integrated eye-blink data in the ASM (N_Categorical_= 43 vs. N_Integrated_= 29) as well as the FPS (N_Categorical_= 55 vs. N_Integrated_= 45) study. Second-level analyses were performed on the estimated parameters for the parametric modulator as calculated within the individual first-levels. A one-sample t-test was performed to find significant associations between neural and muscular activity.

#### Regions of interest and correction for multiple comparisons

For both studies, analyses focused on the CeM and PnC as main regions of interest while BNST (not covered in ASM) and PAG regions were investigated as secondary regions of interest for exploratory purposes.

Based on pre-defined masks for our regions of interest [centromedial amygdala (^64^within the Jülich SPM Anatomy Toolbox (v2.1.) ^65^); BNST ^66^and PAG ^36^], multiple comparisons were controlled for by using a small-volume correction (SVC) approach using family-wise error correction (FWE_SVC_<0.05, cluster-forming threshold at 0.001).

Given that this is the first high-resolution fMRI study targeting the PnC and thus no coordinates as derived from fMRI exist, the brainstem-specific analyses targeting the PnC are reported on a conventional uncorrected threshold of p<0.001. The PnC location was identified by converging information based on anatomical correspondence between structural MRI landmarks and definitions as provided by Duvernoy’s Atlas of the Human Brain Stem and Cerebellum (Naidich et al., 2009, **Figure 1A**). This identified location is additionally supported by its location in reference to a just recently available MRI mask of the nucleus reticularis pontis oralis (PNO, ^36^, **Figure 1A**).

## Code availability

All code used for stimulus presentation, data analyses and figure preparation is available upon request.

## Data availability

FMRI group statistics (T-and F-maps) of all analyses from ASM and FPS presented within the main text are available on Neurovault for download: https://neurovault.org/collections/VLPJMNOM/. Behavioral and psychophysiological data is available upon request.

## References

1. Mobbs, D., Hagan, C. C., Dalgleish, T., Silston, B. & Prévost, C. The ecology of human fear: Survival optimization and the nervous system. Frontiers in Neuroscience 9, (2015).

2. Brown, P. et al. New observations on the normal auditory startle reflex in man. Brain 114, 1891–1902 (1991).

3. Hamm, A. O. in International Encyclopedia of the Social & Behavioral Sciences: Second Edition (2015).

4. Lee, Y., López, D. E., Meloni, E. G. & Davis, M. A primary acoustic startle pathway: obligatory role of cochlear root neurons and the nucleus reticularis pontis caudalis. J. Neurosci. 16, 3775–3789 (1996).

5. Koch, M. The neurobiology of startle. Prog. Neurobiol. 59, 107–128 (1999).

6. Yeomans, J. S. & Frankland, P. W. The acoustic startle reflex: neurons and connections. Brain Res. Rev. 21, 301–314 (1995).

7. Anthony, B. J. in Advances in Psychophysiology (eds. Ackles, P. K., Jennings, J. R. & Coles, M. G. H.) 167–218 (1985).

8. Blumenthal, T. D. et al. Committee report: Guidelines for human startle eyeblink electromyographic studies. Psychophysiology 42, 1–15 (2005).

9. Lang, P. J., Bradley, M. M. & Cuthbert, B. N. Emotion, Attention, and the Startle Reflex. Psychol. Rev. 97, 377–395 (1990).

10. Davis, M., Walker, D. L. & Lee, Y. in Startle modification - Implications for neuroscience, cognitive science, and clinical science (eds. Dawson, M. E., Schell, A. M. & Bohmelt, A. H.) 95–113 (Cambridge University Press, 1999).

11. Grillon, C., Ameli, R., Woods, S. W., Merikangas, K. & Davis, M. Fear-Potentiated Startle in Humans: Effects of Anticipatory Anxiety on the Acoustic Blink Reflex. Psychophysiology 28, 588–595 (1991).

12. Hamm, A. O. & Vaitl, D. Affective learning: Awareness and aversion. Psychophysiology 33, 698–710 (1996).

13. Bradley, M. M., Miccoli, L., Escrig, M. A. & Lang, P. J. The pupil as a measure of emotional arousal and autonomic activation. Psychophysiology 45, 602–607 (2008).

14. Gómez-Nieto, R. et al. Origin and function of short-latency inputs to the neural substrates underlying the acoustic startle reflex. Front. Neurosci. 8, 216 (2014).

15. Hitchcock, J. M. & Davis, M. Lesions of the amygdala, but not of the cerebellum or red nucleus, block conditioned fear as measured with the potentiated startle paradigm. Behav. Neurosci. 100, 11–22 (1986).

16. Hitchcock, J. M. & Davis, M. Efferent pathway of the amygdala involved in conditioned fear as measured with the fear-potentiated startle paradigm. Behav. Neurosci. 105, 826–842 (1991).

17. Rosen, J. B., Hitchcock, J. M., Sananes, C. B., Miserendino, M. J. D. & Davis, M. A Direct Projection From the Central Nucleus of the Amygdala to the Acoustic Startle Pathway: Anterograde and Retrograde Tracing Studies. Behav. Neurosci. (1991).

18. Davis, M. The role of the amygdala in fear-potentiated startle: implications for animal models of anxiety. Trends in Pharmacological Sciences 13, 35–41 (1992).

19. LeDoux, J. E., Iwata, J., Cicchetti, P. & Reis, D. J. Different projections of the central amygdaloid nucleus mediate autonomic and behavioral correlates of conditioned fear. J. Neurosci. 8, 2517–2529 (1988).

20. Davis, M. Neural systems involved in fear and anxiety measured with fear-potentiated startle. Am. Psychol. 61, 741–756 (2006).

21. Davis, M., Walker, D. L. & Lee, Y. Amygdala and bed nucleus of the stria terminalis: differential roles in fear and anxiety measured with the acoustic startle reflex. Philos. Trans. R. Soc. BBiol. Sci. 352, 1675–1687 (1997).

22. Heller, A. S., Lapate, R. C., Mayer, K. E. & Davidson, R. J. The face of negative affect: trial-by-trial corrugator responses to negative pictures are positively associated with amygdala and negatively associated with ventromedial prefrontal cortex activity. J. Cogn. Neurosci. 26, 2102–10 (2014).

23. van Well, S., Visser, R. M., Scholte, H. S. & Kindt, M. Neural substrates of individual differences in human fear learning: evidence from concurrent fMRI, fear-potentiated startle, and US-expectancy data. Cogn. Affect. Behav. Neurosci. 12, 499–512 (2012).

24. Lindner, K. et al. Fear-potentiated startle processing in humans: Parallel fMRI and orbicularis EMG assessment during cue conditioning and extinction. Int. J. Psychophysiol. 98, 535–545 (2015).

25. de Haan, M. I. C. et al. The influence of acoustic startle probes on fear learning in humans. Sci. Rep. 8, 14552 (2018).

26. Wendt, J., Löw, A., Weymar, M., Lotze, M. & Hamm, A. O. Active avoidance and attentive freezing in the face of approaching threat. Neuroimage 158, 196–204 (2017).

27. Schmidt, U., Kaltwasser, S. F. & Wotjak, C. T. Biomarkers in posttraumatic stress disorder: Overview and implications for future research. Disease Markers (2013).

28. Weike, A. I. et al. Fear Conditioning following Unilateral Temporal Lobectomy: Dissociation of Conditioned Startle Potentiation and Autonomic Learning. J. Neurosci. 25, 11117–11124 (2005).

29. Klumpers, F., Morgan, B., Terburg, D., Stein, D. J. & van Honk, J. Impaired acquisition of classically conditioned fear-potentiated startle reflexes in humans with focal bilateral basolateral amygdala damage. Soc. Cogn. Affect. Neurosci. 10, 1161–1168 (2014).

30. Angrilli, A. et al. Startle reflex and emotion modulation impairment after a right amygdala lesion. Brain 119, 1991–2000 (1996).

31. Pissiota, A., Frans, O., Fredrikson, M., Langstrom, B. & Flaten, M. A. The human startle reflex and pons activation: a regional cerebral blood flow study. Eur J Neurosci 15, 395–398 (2002).

32. Pissiota, A. et al. Amygdala and anterior cingulate cortex activation during affective startle modulation: A PET study of fear. Eur. J. Neurosci. 18, 1325– 1331 (2003).

33. Sclocco, R., Beissner, F., Bianciardi, M., Polimeni, J. R. & Napadow, V. Challenges and opportunities for brainstem neuroimaging with ultrahigh field MRI. Neuroimage 168, 412–426 (2018).

34. Beissner, F. Functional MRI of the Brainstem: Common Problems and their Solutions. Clin. Neuroradiol. 25, 251–257 (2015).

35. Naidich, T. P. et al. Duvernoy’s Atlas of the Human Brain Stem and Cerebellum: High-Field MRI, Surface Anatomy, Internal Structure, Vascularization and 3D Sectional Anatomy. (Springer Science & Business Media, 2009).

36. Edlow, B. L. et al. Neuroanatomic connectivity of the human ascending arousal system critical to consciousness and its disorders. J. Neuropathol. Exp. Neurol. (2012).

37. Sabatinelli, D., Lang, P. J., Bradley, M. M., Costa, V. D. & Keil, A. The Timing of Emotional Discrimination in Human Amygdala and Ventral Visual Cortex. (2009).

38. Balaban, M. T. & Taussig, H. N. Salience of fear/threat in the affective modulation of the human startle blink. Biol. Psychol. 38, 117–131 (1994).

39. Dal Monte, O., Costa, V. D., Noble, P. L., Murray, E. A. & Averbeck, B. B. Amygdala lesions in rhesus macaques decrease attention to threat. Nat. Commun. 6, (2015).

40. Dolan, R. J. & Vuilleumier, P. Amygdala Automaticity in Emotional Processing. Ann. N.Y. Acad. Sci. 985, 348–355 (2006).

41. Vuilleumier, P. & Pourtois, G. Distributed and interactive brain mechanisms during emotion face perception: Evidence from functional neuroimaging. Neuropsychologia 45, 174–194 (2007).

42. Wendt, J., Weike, A. I., Lotze, M. & Hamm, A. O. The functional connectivity between amygdala and extrastriate visual cortex activity during emotional picture processing depends on stimulus novelty. Biol. Psychol. 86, 203–209 (2011).

43. Tye, K. M. Neural Circuit Motifs in Valence Processing. Neuron 100, 436–452 (2018).

44. Chekroud, A. M., Lane, C. E. & Ross, D. A. Computational Psychiatry: Embracing Uncertainty and Focusing on Individuals, Not Averages. Biol. Psychiatry (2017).

45. Mumford, J. A., Poline, J. B. & Poldrack, R. A. Orthogonalization of regressors in fMRI models. PLoSOne 10, (2015).

46. Bach, D. R. A cost minimisation and Bayesian inference model predicts startle reflex modulation across species. J. Theor. Biol. 370, 53–60 (2015).

47. Roelofs, K. Freeze for action: Neurobiological mechanisms in animal and human freezing. Philos. Trans. R. Soc. BBiol. Sci. 372, (2017).

48. Glover, E. M. et al. Tools for translational neuroscience: PTSD is associated with heightened fear responses using acoustic startle but not skin conductance measures. Depress. Anxiety 28, 1058–1066 (2011).

49. Hamm, A. O. et al. Panic disorder with agoraphobia from a behavioral neuroscience perspective: Applying the research principles formulated by the Research Domain Criteria (RDoC) initiative. Psychophysiology 53, 312–322 (2016).

50. Wessa, M. et al. EmoPics: Subjektive und psychophysiologische Evaluationen neuen Bildmaterials für die klinisch-bio-psychologische Forschung. Zeitschrift für Klin. Psychol. und Psychother. 1/11, (2010).

51. Brainard, D. H. The Psychophysics Toolbox. Spat. Vis. (1997).

52. Bradley, M. M. & Lang, P. J. Measuring emotion: The self-assessment manikin and the semantic differential. J. Behav. Ther. Exp. Psychiatry 25, 49–59 (1994).

53. Sjouwerman, R., Niehaus, J., Kuhn, M. & Lonsdorf, T. B. Don’t startle me— Interference of startle probe presentations and intermittent ratings with fear acquisition. Psychophysiology 53, 1889–1899 (2016).

54. Mandelkow, H., Halder, P., Boesiger, P. & Brandeis, D. Synchronization facilitates removal of MRI artefacts from concurrent EEG recordings and increases usable bandwidth. Neuroimage 32, 1120–1126 (2006).

55. Haaker, J. et al. Single dose of L-dopa makes extinction memories context-independent and prevents the return of fear. Proc. Natl. Acad. Sci. U.S.A. 110, E2428–36 (2013).

56. Boucsein, W. et al. Publication recommendations for electrodermal measurements. Psychophysiology 49, 1017–1034 (2012).

57. Venables, P. & Christie, M. in Techniques in Psychophysiology 3–67 (Wiley, 1980).

58. Lykken, D. T. & Venables, P. H. Direct measurement of skin conductance: A proposal for standardization. Psychophysiology 8, 656–672 (1971).

59. R Developement Core Team. R: A Language and Environment for Statistical Computing. R Found. Stat. Comput. 1, 409 (2015).

60. Ashburner, J. A fast diffeomorphic image registration algorithm. Neuroimage 38, 95–113 (2007).

61. Diedrichsen, J. A spatially unbiased atlas template of the human cerebellum. Neuroimage 33, 127–138 (2006).

62. Glover, G. H., Li, T. Q. & Ress, D. Image-based method for retrospective correction of physiological motion effects in fMRI: RETROICOR. Magn. Reson. Med. 44, 162–167 (2000).

63. Gläscher, J. Visualization of group inference data in functional neuroimaging. Neuroinformatics 7, 73–82 (2009).

64. Amunts, K. et al. Cytoarchitectonic mapping of the human amygdala, hippocampal region and entorhinal cortex: Intersubject variability and probability maps. in Anatomy and Embryology(2005).

65. Eickhoff, S. B. et al. A new SPM toolbox for combining probabilistic cytoarchitectonic maps and functional imaging data. Neuroimage(2005).

66. Torrisi, S. et al. Resting state connectivity of the bed nucleus of the stria terminalis at ultra-high field. Hum. Brain Mapp. (2015).

