## Supplementary Information for "The neurofunctional basis of affective startle modulation in humans – evidence from combined facial EMG-fMRI"

### Supplementary results

Eye-blink responses in the habituation phase of the ASM study followed a typical trial-by-trial habituation pattern with decreasing response magnitudes over trials. Based on this, parametric modulation analyses revealed two clusters of the CeM (see **Table S1** 'Habituation pattern'**, Figure S1A**) and a cluster in the proximity to the PnC region (**Table S1**) mapping onto the habituation pattern of eye-blink response strength (**Figure 2C**).

**Table S1.** Statistics for PnC activation to startle probe onset during startle probe habituation (unmodulated responding) as well as activation mirroring the EMG habituation pattern in the PnC and secondary regions of interest (CeM, PAG).

| **PnC** | | $p_{uncorrected}$ | **k** | **T** | **X** | **Y** | **Z** |
|---|---|---|---|---|---|---|---|
| *Unmodulated responding* | | | | | | | |
| | | <0.001 | 3 | 3.47 | 2 | -35 | -36 |
| *Habituation pattern* | | | | | | | |
| | | <0.001 | 2 | 3.63 | 6 | -32 | -35 |
| **Secondary ROIs**[a] | | $p_{FWE(SVC)}$ | $k_{(SVC)}$ | **T** | **X** | **Y** | **Z** |
| ***Central Nucleus of the Amygdala*** | | | | | | | |
| *Unmodulated responding* | | | | | | | |
| | left | <0.001 | 80 | 5.23 | -21 | -4 | -14 |
| | right | <0.001 | 24 | 4.90 | 24 | 0 | -15 |
| | | <0.001 | 33 | 4.89 | 20 | -9 | -14 |
| | | <0.001 | | 4.80 | 26 | -9 | -12 |
| | | <0.001 | | 4.65 | 22 | -6 | -12 |
| *Habituation pattern* | | | | | | | |
| | left | 0.008 | 3[b] | 3.75 | 20 | -9 | -14[c] |
| | | 0.013 | 1 | 3.55 | 20 | -4 | -15 |
| ***PAG*** | | | | | | | |
| *Unmodulated responding* | | 0.004 | 13 | 4.16 | -3 | -32 | -10 |
| | | | 9 | 3.58 | 4 | -32 | -10 |

[a] Note, because of the restricted FOV BNST activation was not assessed.
[b] the CeM clusters are part of a larger amygdala cluster with an unrestricted cluster-extend of 120 voxels which lies very specifically within the amygdala boundaries, it might be cautioned to attribute this relationship solely to the area of the CeM. Other areas of the amygdala might also be involved in this habituation related relationship and warrant further investigations.
[c] Note, this is the same peak voxel coordinate as in unmodulated responding.

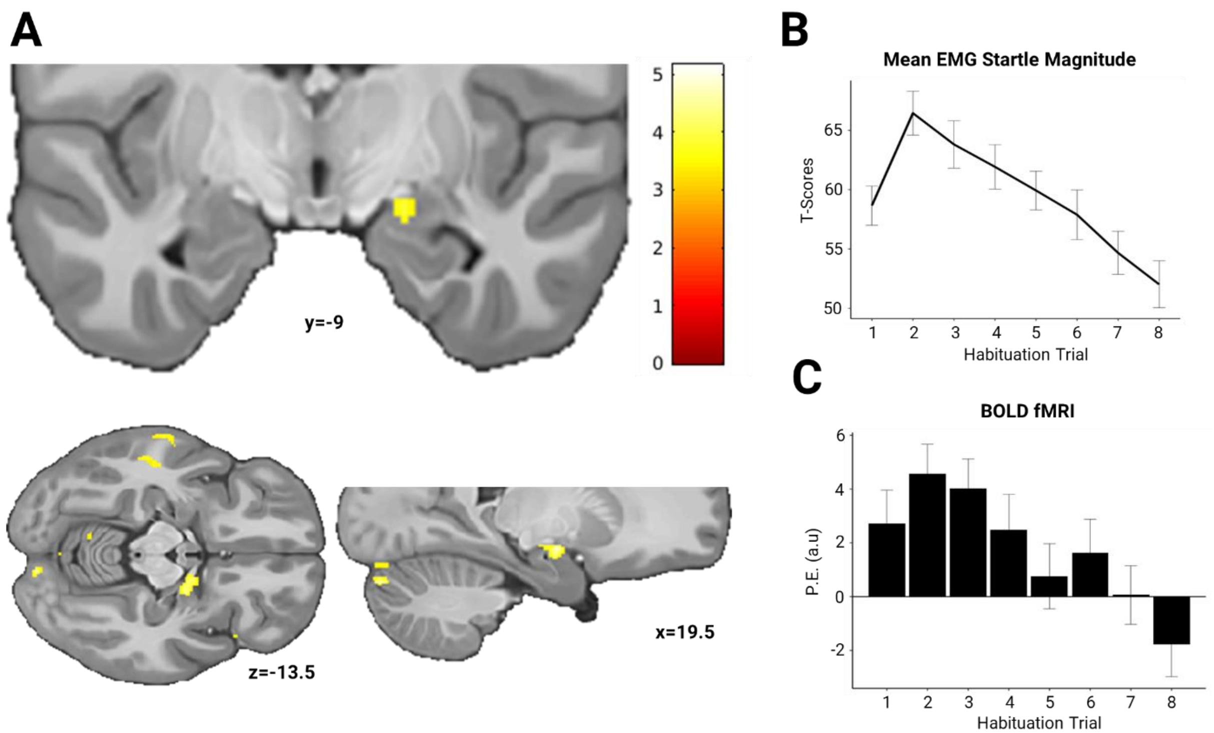


**Figure S1**. (A) Functional mapping of amygdala activation onto the mean trial-by-trial EMG habituation pattern as shown in (B). (C) Extracted estimated parameters from session 1 for the peak amygala voxel in (A) mirroring the EMG pattern in (B). Display threshold at p<0.001. P.E.(a.u.): parameter estimates (arbitrary units).

### Supplementary Discussion

Physiological habituation processes are known to affect PnC activation in the primary acoustic startle reflex in rodents [1]. In line, we observe a characteristic habituation pattern in the PnC with activation strength mirroring EMG magnitude. A similar habituation pattern was observed in the CeM. Importantly, the initial habituation phase may be perceived as unpleasant and the habituation-like pattern may in fact reflect adaptation to the aversive situation. Consequently, adaptation to aversiveness may be considered as a type of affect modulation. It can thus be speculated that the response pattern as observed in the PnC may at least partially reflect affective modulation processes. As such these findings can be taken to support our primary hypotheses presented within the main text.

### References

1. Koch, M. The neurobiology of startle. *Prog. Neurobiol.* **59,** 107–128 (1999).
